# Aberrant splicing in human cancer has a large-scale functional impact on transcription factors

**DOI:** 10.1101/2025.10.16.682783

**Authors:** Khalique Newaz, Olga Tsoy, Jan Baumbach

## Abstract

Cancer shows aberrant alternative splicing (AS), which could functionally perturb proteins required for normal cellular behavior. Transcription factors (TFs) are proteins that regulate gene transcription. Some evidence about AS-driven perturbation of TFs in cancer has been documented. A comprehensive systematic analysis of how cancer-specific AS could affect functions of TFs is missing. Such an analysis could reveal the molecular mechanisms of cancer progression due to transcription misregulations, and thus could help identify therapeutic targets. Here, we systematically analyzed potential functional perturbations of TFs due to AS across 15 cancer types. By analyzing AS patterns, we identified 2 118 perturbed AS events (i.e., events showing significant AS pattern differences between normal and paired cancer samples) that affect 718 TFs across 14 cancer types. In 189 TFs, the perturbed AS events affected known functional domains of TFs. Indirect evidence for the functional impact of cancer-specific AS on TFs were also found: First, by relating AS patterns of the perturbed AS events with the DNA-binding and regulatory activity of TFs. Second, by using cancer dependency data to explore whether the affected TFs are essential for cancer cell line proliferation. Our findings show a large-scale functional perturbation of TFs due to cancer-specific AS.

## 1. Introduction

Alternative splicing (AS) results in multiple different messenger ribonucleic acids (mRNAs) originating from a single gene[1]. AS is associated with normal cellular processes, including tissue development and identity[2]. Normal AS patterns are perturbed in diseases[3], including in cancer[4]. Large-scale RNA sequencing (RNAseq)-based evidence for aberrant AS in cancer has been collected. For example, the cancer genome atlas (TCGA) SpliceSeq database contains RNAseq-based quantification of AS events across 33 cancer types[5]. TCGA SpliceSeq has been used to explore cancer-specific AS[6]. For example, Yongsheg et al.[7] investigated relationships between somatic mutations in cancer and cancer-specific AS patterns, and found that different mutations in the same gene showed distinct AS perturbations. Junyi et al.[8] exploited the hypothesis that RNA-binding proteins (RBPs) regulate cancer-specific AS patterns and proposed a computational pipeline to identify RBPs that are cancer-specific AS regulators. Similarly, Cheng et al.[9] investigated whether abnormal AS of splicing factors could explain cancer-specific AS patterns.

Aberrant AS in cancer has also been shown to impact an important group of proteins called transcription factors (TFs)[10]. TFs bind DNA to regulate transcription of genes[11]. Cancer-specific dysregulation of TFs perturbs normal transcriptional programs, which could lead to a sustained tumor state[12]. Hence, TFs have been considered potential cancer therapeutic targets[13]. TF dysregulations could be due to cancer-specific signaling cascades that perturb TF expression or changes in the corresponding gene structure via chromosomal translocations/point mutations[14]. Some evidence of cancer-specific AS effects on TFs has also been documented[10,15]. For example, AS-induced alternative protein isoforms of T Cell Factor 4 (*TCF4*) have been implicated in proliferations of non-small cell lung cancer cells[16] and hepatocellular carcinoma[17]. Other examples are reviewed elsewhere[10]. Despite existing evidence, a comprehensive systematic analysis of how cancer-specific AS affects TFs has been missing. Such an analysis could reveal how TF-affecting AS events drive cancer progression, which could then be used to identify therapeutic targets and strategize drug design[18].

Here, TCGA SpliceSeq was used to systematically explore the effect of cancer-specific AS events across all known 1 639 human TFs. 15 cancer types that had AS data for enough paired (cancer and healthy) patient samples were used. For each cancer-type, AS events showing statistically significant differences between paired cancer and normal samples were considered as perturbed. 2 118 TF-affecting perturbed AS events covering 718 TFs across 14 cancer types were found. Comparisons across cancer types revealed several unique cancer type-specific, as well as common, TF-affecting perturbed AS events. In 189 of the 718 (∼26%) TFs, at least one perturbed AS event affects TF functional domains. This provides direct evidence that cancer-specific AS events have a functional impact on TFs. Additionally, we found indirect evidence of the functional impact of cancer-specific AS events on TFs in two ways. First, we found that the expression patterns of >100 AS events covering >70 of the 718 TFs have significant correlations with the existing sample-matched TF footprinting data[19], implying a possible dependence of the DNA binding and gene regulatory activities of the affected TFs on the corresponding perturbed AS events. Second, utilizing existing cancer dependency data[20], we found that many (39) TFs affected by perturbed AS events are also essential for cancer progression in cancer cell lines. Overall, our findings highlight a large-scale functional impact of cancer-specific AS on TFs. The collected evidence could guide future experiments to understand cancer molecular mechanisms or identify therapeutic targets.

## 2. Data and methods

### 2.1 Alternative splicing data

From the TCGA database[21] (accessed Feb. 2025), 15 cancer types were selected, where each cancer type had at least 10 patients with primary tumor (cancer) and patient-matched control (healthy) samples (Table 1). For these paired samples, the corresponding AS data were downloaded from TCGA SpliceSeq[5]. This data contains information about seven types of AS events: Alternative Acceptor (AA), Alternative Donor (AD), Alternative Promoter (AP), Alternative Terminator (AT), Exon Skip (ES), Mutually Exclusive Exons (ME), and Retained Intron (RI). To quantify AS events in a TCGA sample, TCGA SpliceSeq provides Percent Spliced In (PSI) values for AS events covered by at least eight RNA-seq reads; for the rest of the AS events, a “null” value is provided. For each cancer type, we only considered events with PSI values for at least 10 paired samples.

**Table 1:** Patient counts across 15 cancer types in TCGA SpliceSeq.

| Cancer type | TCGA cancer code | # of patients with paired samples | # of cancer samples | # of AS events with values for at least 10 paired samples |
| --- | --- | --- | --- | --- |
| Bladder urothelial carcinoma | BLCA | 18 | 406 | 42 343 |
| Breast invasive carcinoma | BRCA | 96 | 1094 | 58 913 |
| Colon adenocarcinoma | COAD | 39 | 458 | 48 843 |
| Esophageal carcinoma | ESCA | 13 | 180 | 44 778 |
| Head and neck squamous cell carcinoma | HNSC | 42 | 501 | 51 442 |
| Kidney chromophobe | KICH | 25 | 66 | 50 059 |
| Kidney renal clear cell carcinoma | KIRC | 71 | 533 | 57 286 |
| Kidney renal papillary cell carcinoma | KIRP | 32 | 290 | 52 337 |
| Liver hepatocellular carcinoma | LIHC | 50 | 371 | 43 753 |
| Lung adenocarcinoma | LUAD | 51 | 514 | 51 436 |
| Lung squamous cell carcinoma | LUSC | 48 | 501 | 57 040 |
| Prostate adenocarcinoma | PRAD | 51 | 497 | 55 799 |
| Stomach adenocarcinoma | STAD | 33 | 415 | 54 446 |
| Thyroid carcinoma | THCA | 57 | 505 | 56 824 |
| Uterine corpus endometrial carcinoma | UCEC | 23 | 545 | 36 585 |

For each cancer type, significantly perturbed AS events were detected as follows. For an AS event, only the patients with PSI values for both normal and cancer samples were detected. Considering these patients, a two-sided paired Wilcoxon Signed Rank test was performed to compare PSI values from normal samples and the paired cancer samples using the “*wilcox.test*” function in *R* to obtain a p-value. The p-values across all AS events were corrected using the Benjamini Hochberg procedure to obtain the corresponding q-values. An AS event was considered as significantly perturbed if and only if (1) its q-value was < 0.05 and (2) the difference between the means of normal and cancer PSI values was >5%.

### 2.2 Transcription factors

#### 2.2.1 Curation and mapping of genomic coordinates

All 1 639 curated human TFs were obtained from Supplementary Table S1 of the Lambert et al.[11] study. For these TFs, canonical protein sequences from UniProt were collected. Henceforth, by the term “UniProt protein sequence” we mean the “canonical UniProt protein sequence”. UniProt IDs and Ensembl transcripts were matched using UniProt. Because one TF can have multiple Ensembl transcripts, one transcript per TF was selected as follows. Given a TF, its UniProt sequence was pairwise globally aligned with each of the corresponding Ensembl transcripts. Protein sequences of the transcripts were constructed using the release 113 of Genome Reference Consortium Human Build 37 (GRCh37) Ensembl GTF file “Homo_sapiens.GRCh37.dna.primary_assembly.fa.gz”. Then, the transcript that matched the TF protein sequence with 100% identity and no gaps was selected. For the 1 469 TFs with matched transcripts as per the above criterion, genomic coordinates were mapped onto their protein sequences using GRCh37. Note that GRCh37 was used instead of GRCh38 because TCGA SpliceSeq used GRCh37 to map genomic coordinates of the AS events. Genomic coordinates of AS events provided by TCGA SpliceSeq were then used to mark parts of the TF protein sequences affected by AS events.

Specifically, for each of the six types of AS events, i.e., AA, AD, AP, AT, ES, and ME, the following was done. Note that RI AS events can not be mapped to TFs as they do not overlap with annotated protein-coding regions defined by GRCh37. For AA, AD, ES, or ME, all sequence positions of a TF whose start and stop genomic coordinates lie within the start and stop coordinates of the affected exons of an AS event were marked as affected. For AP, all sequence positions of a TF whose start genomic coordinates lie before the start position of the affected exon were marked as affected. For AT, all sequence positions of a TF whose stop genomic coordinates lie beyond the stop position of the affected exon were marked as affected.

**Figure 1.**
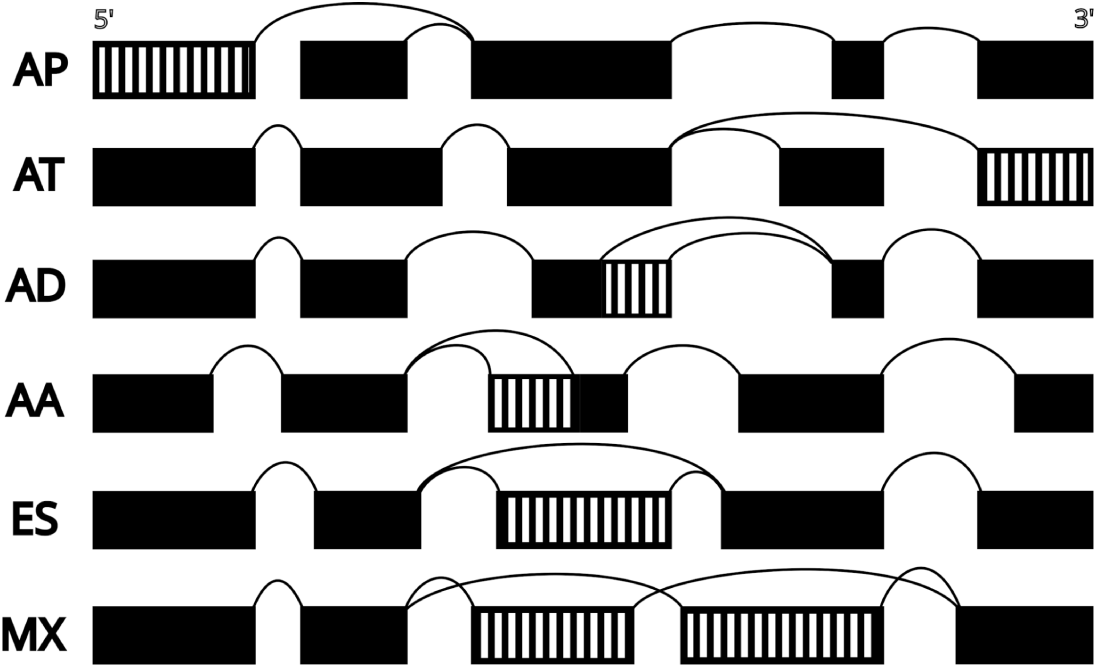
Schematic illustration of AS affected exons (marked as striped boxes). 1st row: for the AS event using the second exon as AP, the first exon was marked as affected. 2nd row: for the AS event using the second last exon as AT, the last exon was marked as affected. 3rd row: for the AS event using an alternative 3’ end of an exon (AD), the part of the exon beyond alternative 3’ was marked as affected. 4th row: for the AS event using an alternative 5’ end of an exon (AA), the part of the exon before alternative 5’ was marked as affected. 5th row: for the AS event skipping an exon (ES), the corresponding exon was marked as affected. 6th row: for the AS event involving two mutually exclusive exons (MX), both exons were marked as affected.

#### 2.2.2 Mapping of DNA-binding and effector domains

UniProt collects DNA-binding domain (DBD) information of proteins using PROSITE[22], Pfam[23], or SMART[24]. This DBD information was used to annotate which sequence positions of TFs belong to a DBD region. Specifically, given the information about a TF in UniProt, if present, the following feature table entries were considered as a DBD: (1) entries “DNA_BIND” or “ZN_FING” or (2) entry “DOMAIN” with domain names ‘bHLH’, ‘bZIP’, or ‘MADS-box’. Of the 1 469 TFs with a matching Ensembl transcript (Section 2.2.1), 1 347 had at least one sequence position belonging to a DBD. TF effector domain (ED) information was taken from Supplementary Tables S2 and S3 of the Soto et al.[25] study. This study looked for ED evidence across 1 639 human TFs in the literature and manually curated the largest collections of 924 EDs across 594 TFs. For these TFs information about the UniProt sequence (the corresponding isoform) positions of EDs are provided. This ED information was used to annotate which UniProt sequence positions of TFs belong to an ED region. Of the 1 469 TFs (Section 2.2.1), 460 had at least one sequence position belonging to an ED region, covering 730 EDs.

### 2.3 tSNE application

Across all 15 cancer types, 2 278 TF AS events were identified that were present in each cancer type. Among these, 1 011 TF AS events were present as perturbed TF splicing events (PTSEs) in at least one cancer type. PSI patterns of these 1 011 AS events for all 6 876 cancer samples across the 15 cancer types were used as input to the “*Rtsne*” package of R to obtain dimensionally reduced representations of the cancer samples.

### 2.4 Hypergeometric test of DBD perturbations

Among the 1 347 TFs with DBD annotation (Section 2.2.2), 1 464 occurrences of DBDs were found, which were taken as the background size. Among all PTSEs across all cancer types, 146 unique DBDs were present, which were taken as the sample size. For each of the 27 DBD-types perturbed in at least one PTSE, their counts in the sample and the background were used to compute the corresponding over-representations via the hypergeometric test to obtain 27 p-values. Similarly, for each of the 27 DBD-types, their counts in the sample and the background were used to compute the corresponding under-representations via the hypergeometric test to obtain 27 p-values. The 54 hypergeometric p-values were corrected using the Benjamini Hochberg procedure to obtain the corresponding q-values.

### 2.5 Alternative splicing events significantly associated with patient survival

Associations between AS events from TCGA SpliceSeq and cancer patient survival were obtained from the OncoSplicing database (“SpliceSeq_info_Survival.csv.gz”) developed in Zhang et al.[26]. For a cancer type, OncoSplicing contains AS events that correlate with clinical outcomes according to the Cox proportional hazards regression. All AS events with survival association p-values of less than 0.05 (reported in the column named ‘Pvalue_HR’ of “SpliceSeq_info_Survival.csv.gz”) were considered as significant and selected for our study.

### 2.6 Correlation of PSI values of PTSEs and TF footprinting

TF footprinting data were taken from Supplementary Table S6 of Corces et al.[19], which covers 410 TCGA cancer samples and 843 TFs across 23 cancer types. Corces et al. provides footprinting depth and flanking accessibility values for 1 781 TF-DNA motif pairs, with multiple DNA motifs for a given TF. For a cancer type, a correlation between PSI values of a PTSE and footprinting values of a TF-DNA motif pair was measured if and only if there were at least 10 cancer samples for which the PTSE had a non-null PSI value in TCGA SpliceSeq and footprinting data in Corces et al. At least 10 cancer samples were considered to retain power in the correlation analysis. 769 PTSEs across 12 cancer types fulfilled this criterion (Table 2).

**Table 2:** Overlap between TCGA SpliceSeq and Corces et al.^19^.

| TCGA<br>cancer code | # of cancer<br>samples | # of PTSEs with PSI values for<br>at least 10 cancer samples | # of TFs within PTSEs with PSI values<br>for at least 10 cancer samples |
| --- | --- | --- | --- |
| BLCA | 10 | 42 | 28 |
| BRCA | 74 | 296 | 132 |
| COAD | 38 | 176 | 88 |
| KIRC | 16 | 245 | 116 |
| KIRP | 33 | 150 | 84 |
| LIHC | 17 | 93 | 37 |
| LUAD | 20 | 198 | 93 |
| LUSC | 16 | 299 | 135 |
| PRAD | 26 | 124 | 67 |
| STAD | 21 | 96 | 56 |
| THCA | 14 | 144 | 78 |
| UCEC | 13 | 85 | 52 |

Given a cancer type, for each PTSE selected as per the above criterion, we measured the correlation between its PSI values and footprinting depth of the corresponding TF-DNA motif pairs. Over all 12 cancer types this resulted in 3 676 correlation pairs. Similarly, we obtained 3 676 correlation pairs corresponding to flanking accessibility. We calculated the empirical p-value based on the data randomization. For each of the 3 676*2 = 7 352 pairs, 100 random correlations were computed by randomly shuffling the PSI values and the footprinting data. The fraction of random correlations higher than the real correlation was taken as the empirical p-value. The resulting 7 352 p-values were corrected using the Benjamini Hochberg procedure to obtain the corresponding q-values. A q-value < 0.05 was considered significant.

### 2.7 Dependency profiles of TFs associated with PTSEs

Dependency profiles of cancer cell lines were downloaded from the depmap[20] portal (“https://depmap.org/portal/data_page/?tab=customDownloads”), which included all available cell lines and genes. This data cover 12 cancer types considered in our study: BLCA, BRCA, COAD, HNSC, KIRC, LIHC, LUAD, LUSC, PRAD, STAD, THCA, and UCEC. For these 12 cancer types, cell lines corresponding the following terminologies (in the order of the cancer types mentioned above) from the “lineage_3” column of the data was used: “Bladder Urothelial”, “Breast Invasive Carcinoma”, “Colorectal Adenocarcinoma”, “Head and Neck Squamous Cell Carcinoma”, “Renal Clear Cell Carcinoma”, “Hepatocellular Carcinoma”, “Lung Adenocarcinoma”, “Lung Squamous Cell Carcinoma”, “Prostate Adenocarcinoma”, “Stomach Adenocarcinoma”, “Thyroid Cancer”, “Endometrial Carcinoma”. The PTSEs across all of the above 12 cancer types cover 688 TFs. Of the 688 TFs, for 669 TFs across the 12 cancer types, there is dependency data with respect to at least one cancer cell line, totaling 2 505 TF-cancer type combinations.

## 3. Results and discussion

### 3.1 Perturbed TF splicing events in individual cancer types

Given a cancer type, all AS events showing statistically significant (q-value < 0.05 and a difference of >5% between the means of PSI values from normal and cancer samples) PSI differences were considered as perturbed AS events (Section 2.1). From these, all AS events corresponding to TFs (Section 2.2) were considered as perturbed TF splicing events (PTSEs).

Over all cancer types, 718 TFs were affected by at least one PTSE. Many PTSEs were found for each cancer type, except for ESCA that showed no perturbed AS event and thus no PTSE. No perturbed AS event for ESCA could be due to the low number of samples (13; see Table 1) that do not provide enough statistical power to detect significant differences between cancer and normal samples with the current q-value cutoff. The number of PTSEs showed large variations across cancer types, with the highest number (750) of PTSEs for LUSC (Figure 2A). On average over all cancer types, ∼1.8 PTSEs came from every TF, implying that no TF dominates PTSEs. PTSE numbers within cancer types do not seem to correlate with the corresponding numbers of paired normal and cancer samples (Table 1). For example, LIHC, LUAD, LUSC, and PRAD, had similar numbers (50, 51, 48, and 51 respectively) of paired samples, but their PTSE numbers varied substantially: 247, 461, 750, and 310, respectively. Overall, these results indicate that not only are there significant TF AS differences between normal and cancer samples within a cancer type, but these differences also vary considerably across cancer types.

**Figure 2.**
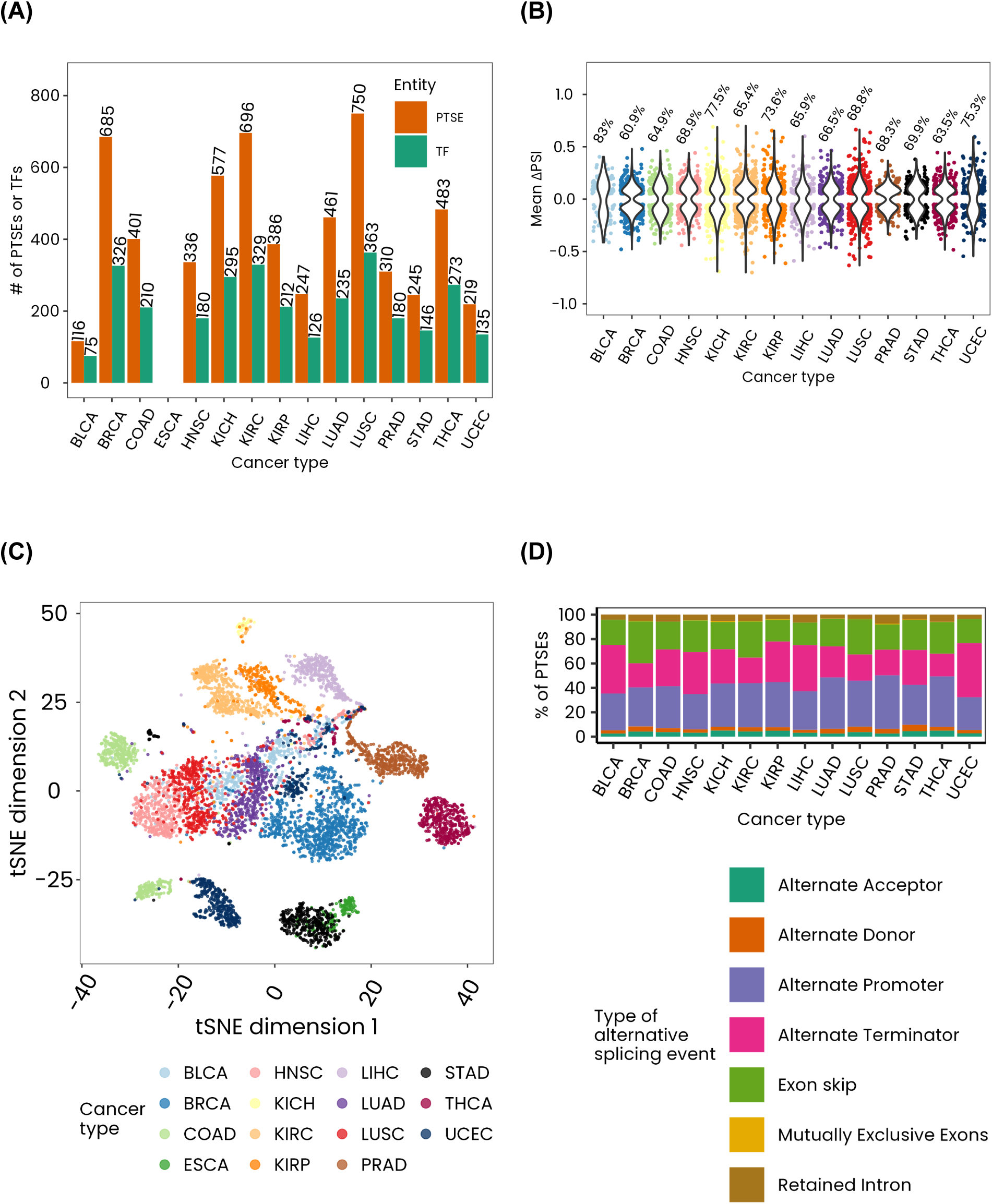
Cancer type related PTSEs. **(A)** Number of cancer type related PTSEs. **(B)** Distribution of mean ΔPSI values of PTSEs. For a cancer type, the y-coordinate of a point (PTSE) represents the ΔPSI averaged over all patients with paired samples. The top of a violin bar represents the average percentage of patients in which a PTSE is consistently higher/lower in cancer samples than in paired normal samples. For example, for BLCA, this percentage is 83%. **(C)** t-SNE visualization of the cancer samples of the entire TCGA cohort. **(D)** Percentage distribution of the types of AS events.

For many PTSEs within a cancer type, their PSI values of cancer samples were higher/lower than their PSI values of the paired normal samples for a majority of patients (Figure 2B). For example, the PSI values for the AS event *HMGA2_22880_AT* (here, “HMGA2” is the gene symbol, “22880” is the AS event number provided by the TCGASpliceSeq database, and “AT” is the type of AS event) were higher in cancer than normal samples for 52 out of the 57 paired samples of THCA with a mean ΔPSI of around 0.44 (Supplementary Table S1). Here, ΔPSI was defined as the difference between the PSI value of a cancer sample and the paired normal sample, with mean ΔPSI denoting the average ΔPSI over all paired samples of a given cancer type. The above results imply cancer type relevance of PTSEs. To further check whether this is the case, we used t-distributed Stochastic Neighbor Embedding (tSNE) to visualize cancer samples of the entire cohort of the considered cancer types, based on PSI patterns of all 1 011 AS events that were present in each of the 15 cancer types and were present as PTSEs in at least one cancer type (Section 2.3). We found that cancer type-related samples were generally clustered together (Figure 2C), implying cancer tissue specificity of the PSI patterns of PTSEs. Similar results have been shown using PSI patterns of all AS events in cancer[8,9]. Interestingly, although we found large variations in PTSEs with respect to their numbers and PSI patterns across cancer types, the proportions of different types of AS events within PTSEs were similar across cancer types. AP, AT, and ES dominate PTSEs (Figure 2D), which has also been found in existing studies using all AS events[8,9].

### 3.2 Common and unique PTSEs of individual cancer types

Of the total 2 118 PTSEs, a majority (∼61%, i.e., 1 302 PTSEs) occurred in more than one cancer type (Figure 3A and Supplementary Table S2). 39 PTSEs occurred in at least 10 cancer types (Figure 3B). An example PTSE is *PHD19_87401_AT* that affects the PHD Finger Protein 19 (*PHD19*) gene. This PTSE was upregulated in cancer samples in 12 cancer types, and corresponds to a known PHF19 isoform (hPCL3s) that is related to cancer progression of multiple cancer types[27–30]. We also found many unique PTSEs (816 in total) for individual cancer types that varied across cancer types, with BRCA having the most (151) and BLCA having the least (6) of them (Figure 3C). Interestingly, at least one of the unique PTSE is survival-associated with the corresponding cancer type (Section 2.5), but not to any of the other cancer types (Figure 3C and Supplementary Table S3). For example, PTSE *PITX2_70356_AP* affects the Pituitary homeobox 2 (*PITX2*) gene that results in the PITX2 isoform named PITX2D[31]. *PITX2_70356_AP* was upregulated in cancer samples compared to normal samples in BLCA with a mean ΔPSI of 0.27, and was found to be associated with disease-free interval of BLCA with a hazard ratio of 0.31 (Supplementary Table S3). This means that an increase in the inclusion of this PTSE indicates a decreased risk of BLCA recurrence. PITX2 isoforms have been shown to be crucial for developmental processes, with PITX2D inhibiting the transcriptional activities of two other PITX2 isoforms by physically interacting with them[31]. This might explain why the increased expression of *PITX2_70356_AP*, which potentially leads to an increased expression of PITX2D, correlates with a decrease in the risk of BLCA recurrence. These results suggest that unique PTSEs could be relevant to understand the molecular mechanisms of disease progression of individual cancer types.

**Figure 3.**
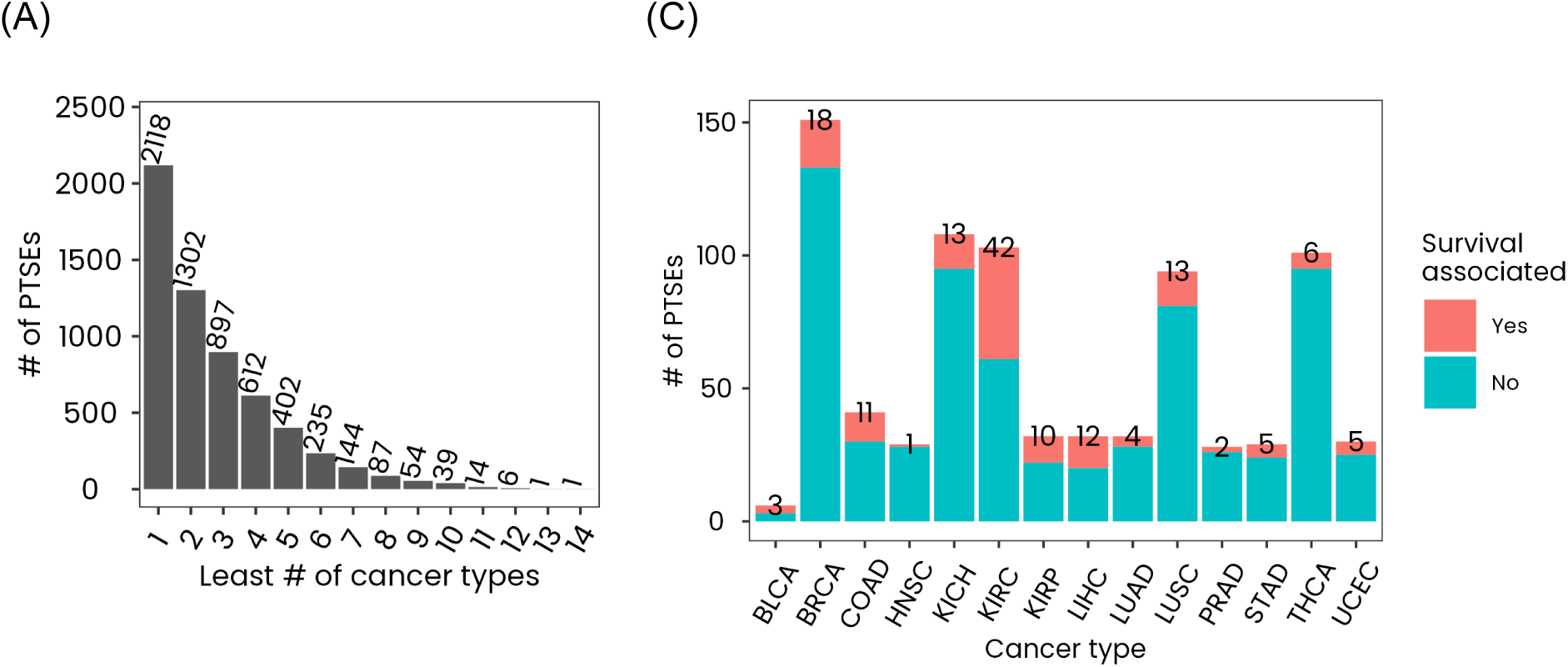
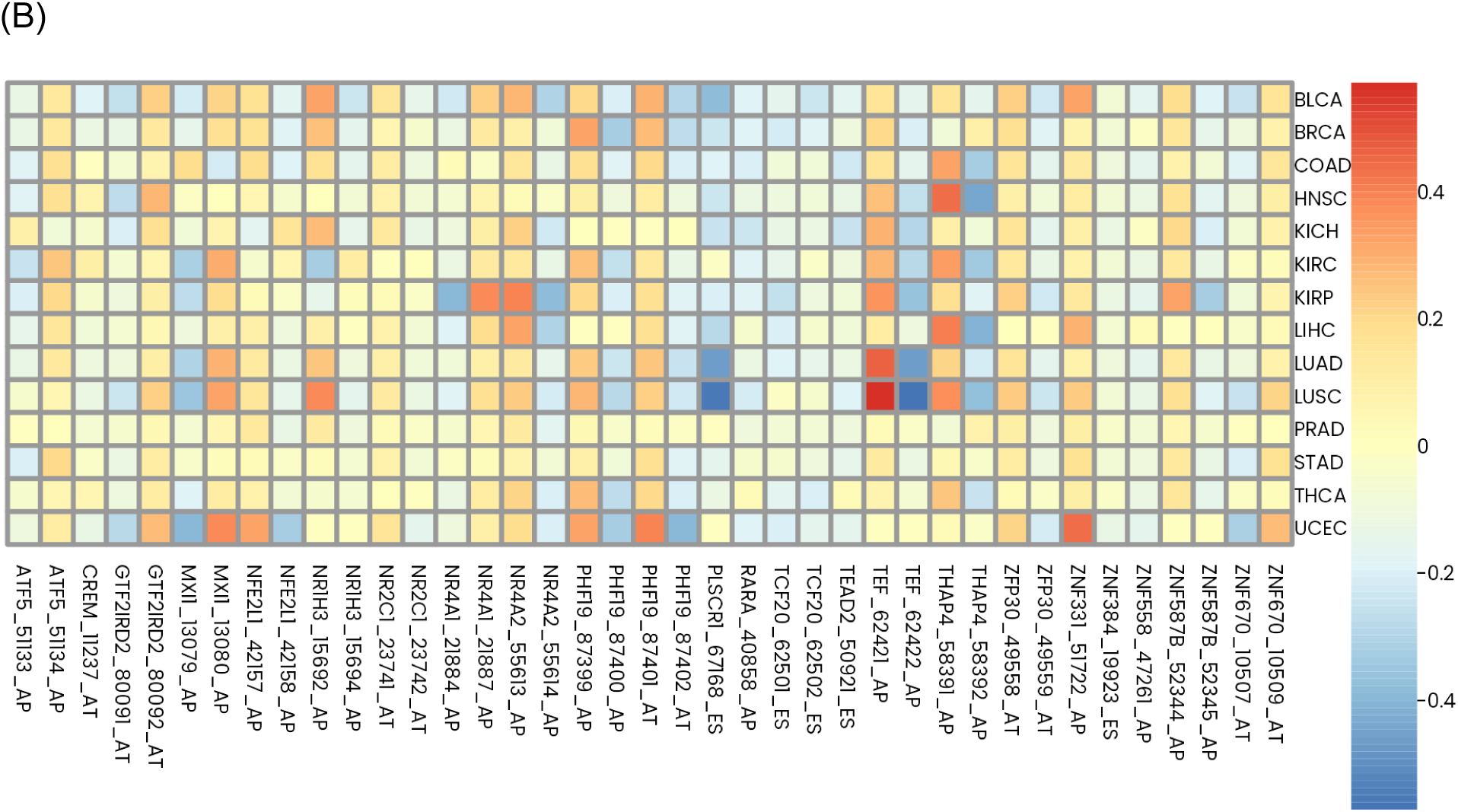
Common and unique PTSEs. **(A)** Number of PTSEs that occur in one or more cancer types. **(B)** Mean ΔPSI of the 39 PTSEs (y-axis) present in at least 10 cancer types (x-axis). **(C)** Number of unique PTSEs of individual cancer types, with the number of such PTSEs that are uniquely associated with patient survival denoted on top of the bars. Information about survival association was derived from Zhang et al.[26].

### 3.3 PTSEs perturb DNA-binding domains

DBDs are important functional components of TFs, which enable TFs to recognize and bind the specific DNA sequences related to their target genes[11]. AS events affecting DBDs can impact the DNA-binding ability of the resulting TF isoforms. To capture this, we evaluated whether PTSEs affect DBD regions of TFs (Section 2.2).

Over all cancer types, DBD regions of 135 TFs (i.e., ∼10% of all 1 347 TFs with a DBD information; Section 2.2) were perturbed by at least one PTSE (Supplementary Table S4). Similar to the numbers of all PTSEs (Figure 3A), numbers of DBD perturbing PTSEs show large variations across cancer types, with the highest number (87) for LUSC and the lowest number (8) for BLCA (Figure 4A). Many of such PTSEs affect large portions of DBD regions within protein sequences of TFs (Figure 4B), and thus could have a strong functional impact. An interesting example is the PTSE *ELF5_14951_AT*, which perturbs the Erythroblast Transformation Specific (ETS) domain of E74-like transcription factor 5 (ELF5). This PTSE has higher PSI values in cancer samples than in normal samples of kidney cancer: KICH, KIRP, and KIRC (Figure 4B). *ELF5_14951_AT* seldom shows large differences among normal vs. cancer samples in other cancer types (Supplementary Table S1), implying its potential functional relevance in kidney cancer. Interestingly, ELF5 has been shown to be a lineage-specific TF of principal cells that are key cells involved in kidney development[32]. Overall, these results indicate that PTSEs could have functional relevance in cancer phenotype development.

**Figure 4.**
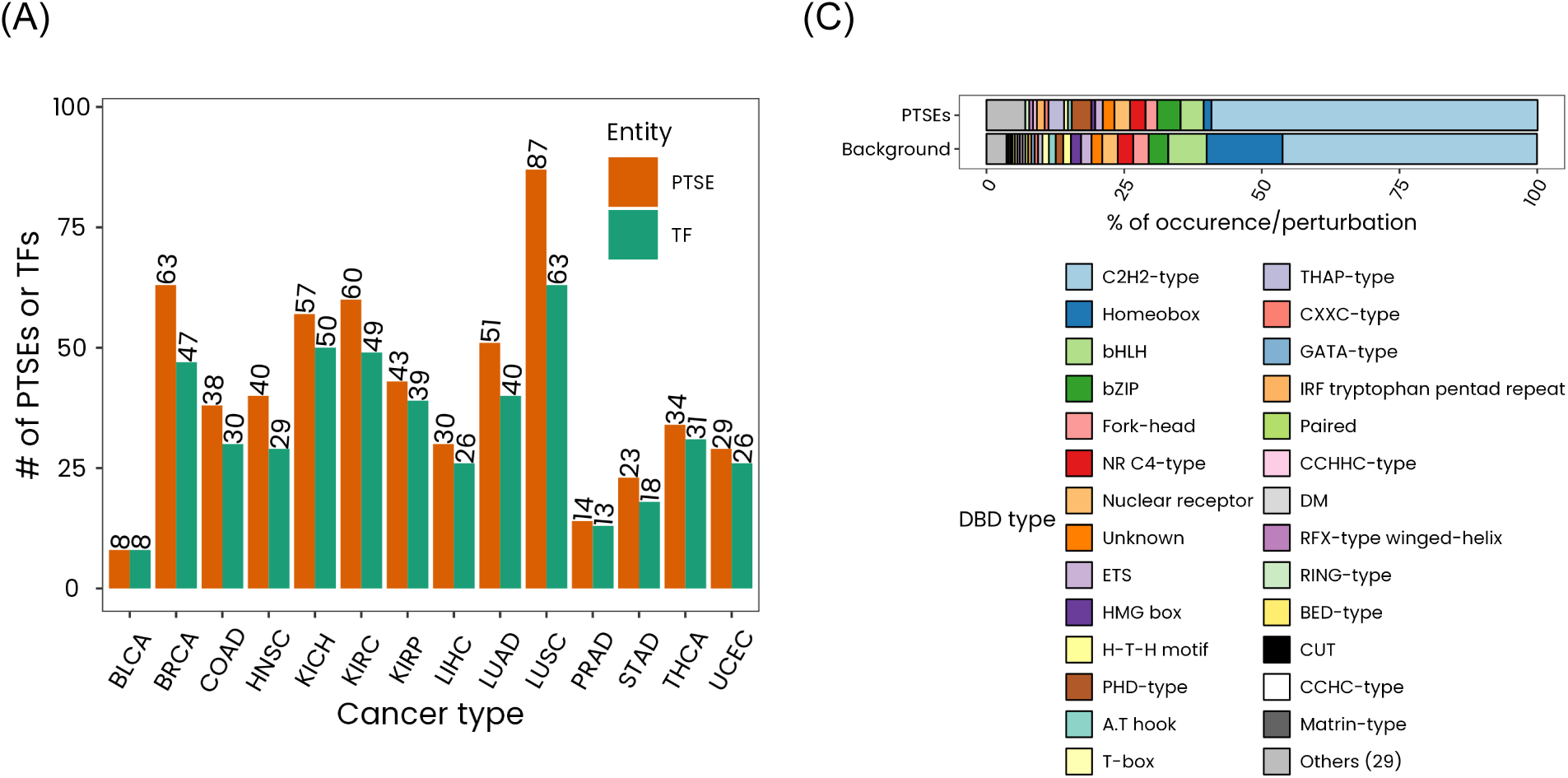
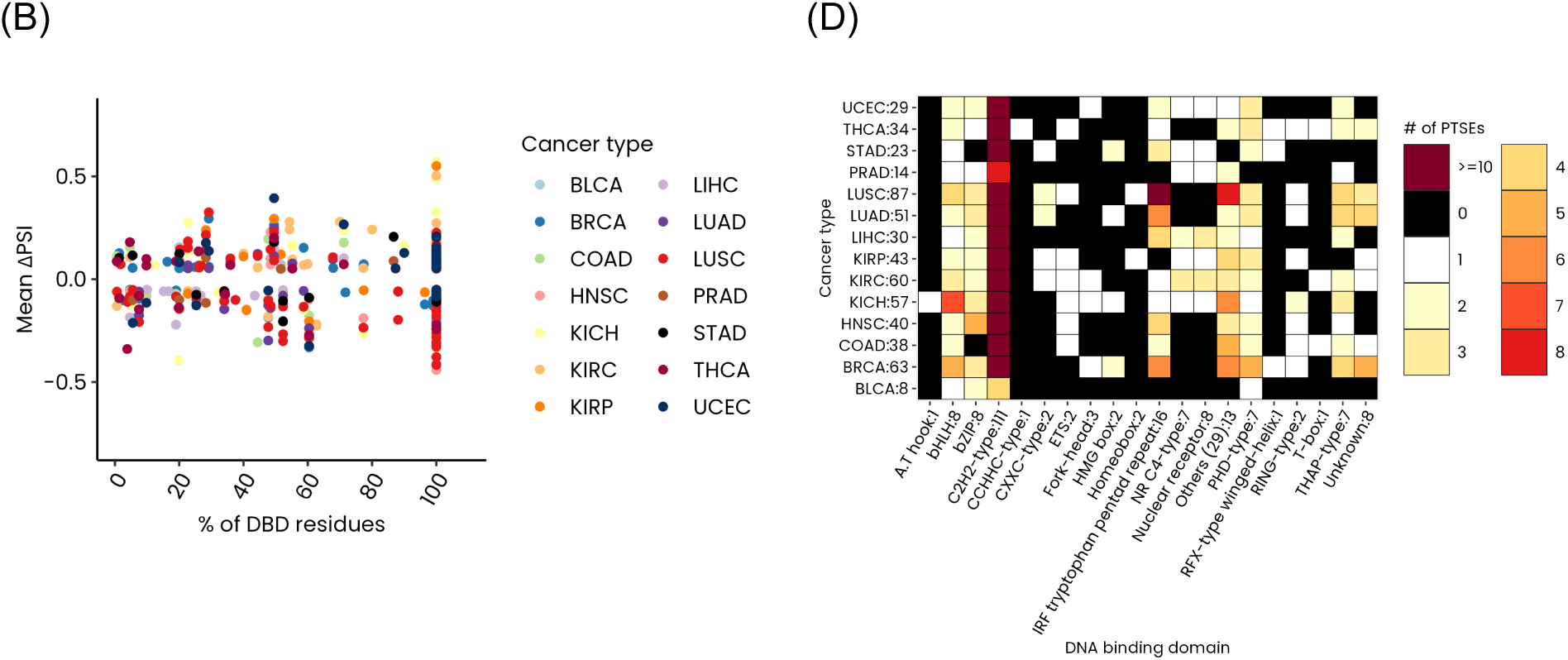
PTSEs that perturb DBDs. (A) Number of cancer type related PTSEs that perturb DBDs. (B) Distribution of the percentage of DBD residues perturbed by PTSEs. Each point represents a PTSE (with the corresponding mean ΔPSI on the y-axis) that results in a DBD perturbation (with the percentage of the perturbed DBD region on the x-axis). (C) Percentage distribution of the DBD types within all considered TFs (background) and those within PTSEs. For visualization purposes, 29 DBD types with counts of less than 5 in the background set of 1 347 TFs were grouped into one as “Others (29)”. (D) Perturbed DBD types within PTSEs. The x-axis shows the names of DBD types and the corresponding number of PTSEs separated by colon. For example, for “bHLH:8” on the second column from left denotes that bHLH is perturbed in 8 PTSEs. The y-axis shows the cancer types and the corresponding number of PTSEs separated by colon. For example, for “BLCA:8” on the first row from bottom, denotes that BLCA has 8 PTSEs.

We next evaluated whether certain DBD types are significantly over/under-represented within PTSEs. Specifically, we quantified the number of DBDs within PTSEs of any cancer type compared to the number of DBDs over all considered TFs, using the hypergeometric test (Section 2.4). The proportions of DBDs among all considered TFs and those present in PTSEs were similar, with one significant exception (Figure 4C). That is, the Homeobox domain was significantly under-represented (q-value < 10^-06^) in PTSEs. Under-representation of the Homeobox domain in cancer was also observed in a recent study that detected 1 350 TF-DNA interactions among 265 TFs and cloned promoters of 108 cancer-related genes[33]. These results mean that the Homeobox domain is not only less relevant for cancer-related gene regulations in general, but it is also less affected by cancer-related AS events. Within individual cancer types, the DBD types show different perturbation patterns (Figure 4D). Some DBD types (i.e., bHLH, bZIP, C2H2-type, IRF tryptophan pentad repeat, PHD-type, and THAP-type) were perturbed in multiple (>12) cancer types, while others (i.e., A.T hook, CCHHC-type, and Homeobox) were perturbed in selected (<3) cancer types. These results could reflect the underlying AS-induced shared and unique TF-DNA binding perturbations among cancer types.

### 3.4 PTSEs perturb effector domains

Besides DBDs, effector domains (EDs) also constitute important functional components of TFs that enable activation or repression of the target genes[25]. Thus, AS events affecting EDs can also impact the functionality of the resulting TF isoforms. To capture this, we evaluated whether PTSEs affect ED regions of TFs (Section 2.2).

Similar to the numbers of DBD perturbing PTSEs (Figure 4A), numbers of ED perturbing PTSEs show large variations across cancer types, with the highest number (51) for KIRC and the lowest number (14) for BLCA (Figure 5A and Supplementary Table S5). Similar to DBD perturbing PTSEs, many ED perturbing PTSEs affect large portions of ED regions within TF protein sequences (Figure 4B). For example, the PTSE *TP73_321_AP* perturbs the activator domain (AD) of TP73 in LUSC. This PTSE has higher PSI values in cancer samples than in normal samples of LUSC (Figure 5B). *TP73_321_AP* leads to the ΔNp73 isoform of TP73, which is known to be overexpressed in tumors compared to normal conditions[34] and has been shown to fail in inducing apoptosis and cell cycle arrest[35].

**Figure 5.**
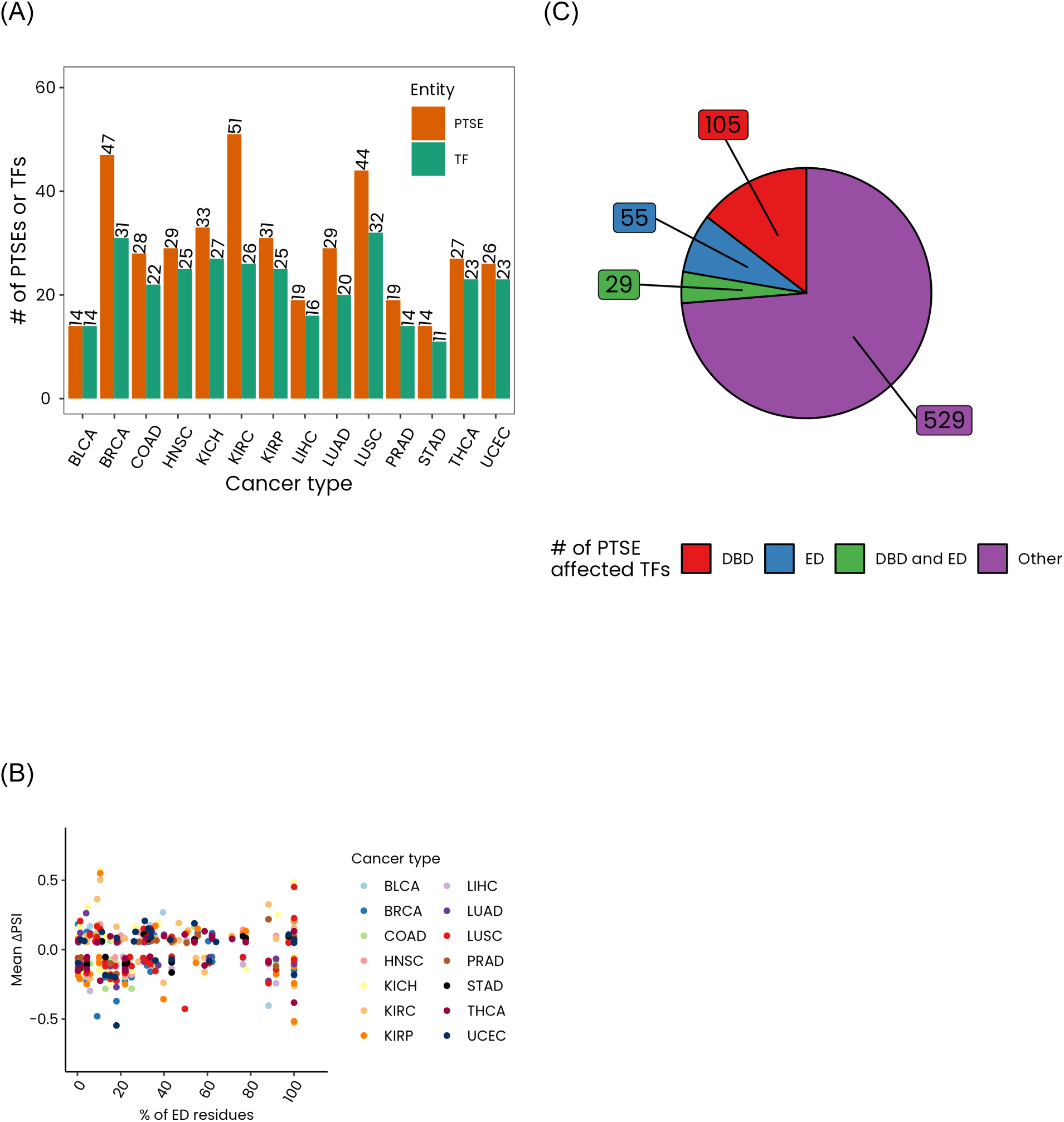
PTSEs that perturb EDs. **(A)** Number of cancer type related PTSEs that perturb EDs. **(B)** Distribution of the percentage of ED residues perturbed by PTSEs. Each point represents a PTSE (with the corresponding delta PSI on the y-axis) that results in an ED perturbation (with the percentage of the perturbed ED on the x-axis). **(C)** Number of TFs with PTSE-induced perturbations in their DBD regions, ED regions, both DBD and ED regions, or other protein regions.

Over all cancer types, 149 PTSEs perturb ED regions across 84 TFs (i.e., ∼18% of all 460 TFs with ED information; Section 2.2). This proportion is almost twice the proportion of TFs (∼10%) whose DBD regions were perturbed by PTSEs. This implies that PTSEs more frequently affect the transcriptional activity of TFs than their DNA binding ability. Of all 718 TFs that are perturbed by at least one PTSE, 189 (∼26%) involve a DBD or an ED, of which 29 TFs show perturbations in both ED and DBD regions. These results imply that AS causes large-scale functional perturbation of TFs in cancer.

### 3.5 Alternative splicing patterns of PTSEs correlate with TF footprinting

To determine potentially AS-driven TF binding/activity patterns, we correlated the PSI values of PTSEs with the TF footprinting of the corresponding TFs in cancer samples of individual cancer types (Section 2.6). To do this, the chromatin accessibility data from Corces et al.[19] was used. Corces et al. used assay for transposase-accessible chromatin using sequencing (ATAC-seq) to determine chromatin occupancy of TFs in TCGA cancer samples, quantified via “footprinting depth” and “flanking accessibility”. Intuitively, footprinting depth measures the inverse of the extent to which the binding of a TF to a chromatin region protects the corresponding TF-binding DNA motif from DNase[36]. Thus, low footprinting depth means high TF binding, and vice versa. In contrast, flanking accessibility measures the accessibility of the DNA region adjacent to TF-binding DNA motifs, which has been associated with TF activity[37]. Thus, high flanking accessibility could mean high TF activity, and vice versa.

We found 122 PTSEs covering 77 TFs across 11 cancer types (146 unique PTSE-cancer type combinations) that showed significant (q-value < 0.05) correlation with footprinting depth, flanking accessibility, or both (Figure 6 and Supplementary Table S6). Many of these PTSEs showed high mean ΔPSI values. For example, the PTSE *PPARG_63413_AP* affecting Peroxisome Proliferator-Activated Receptor Gamma (PPARG) in LUSC is significantly negatively correlated with footprinting depth and positively correlated with flanking accessibility. This could imply that, in LUSC, with an increase in the proportion of *PPARG_63413_AP* there is an increase in the DNA binding as well as the gene regulatory activity of PPARG. Because *PPARG_63413_AP* shows a mean ΔPSI of −0.42 in LUSC, DNA binding and gene regulatory activities of PPARG is perhaps higher in the normal tissue than in LUSC. Interestingly, previous in vivo[38] and in vitro[39] studies have shown that an increase in the PPARG activity slows down the tumor growth rate in non-small-cell lung cancer. Overall, this analysis identified PTSEs within cancer types whose inclusion rates are correlated with the activities of the corresponding TFs.

**Figure 6.**
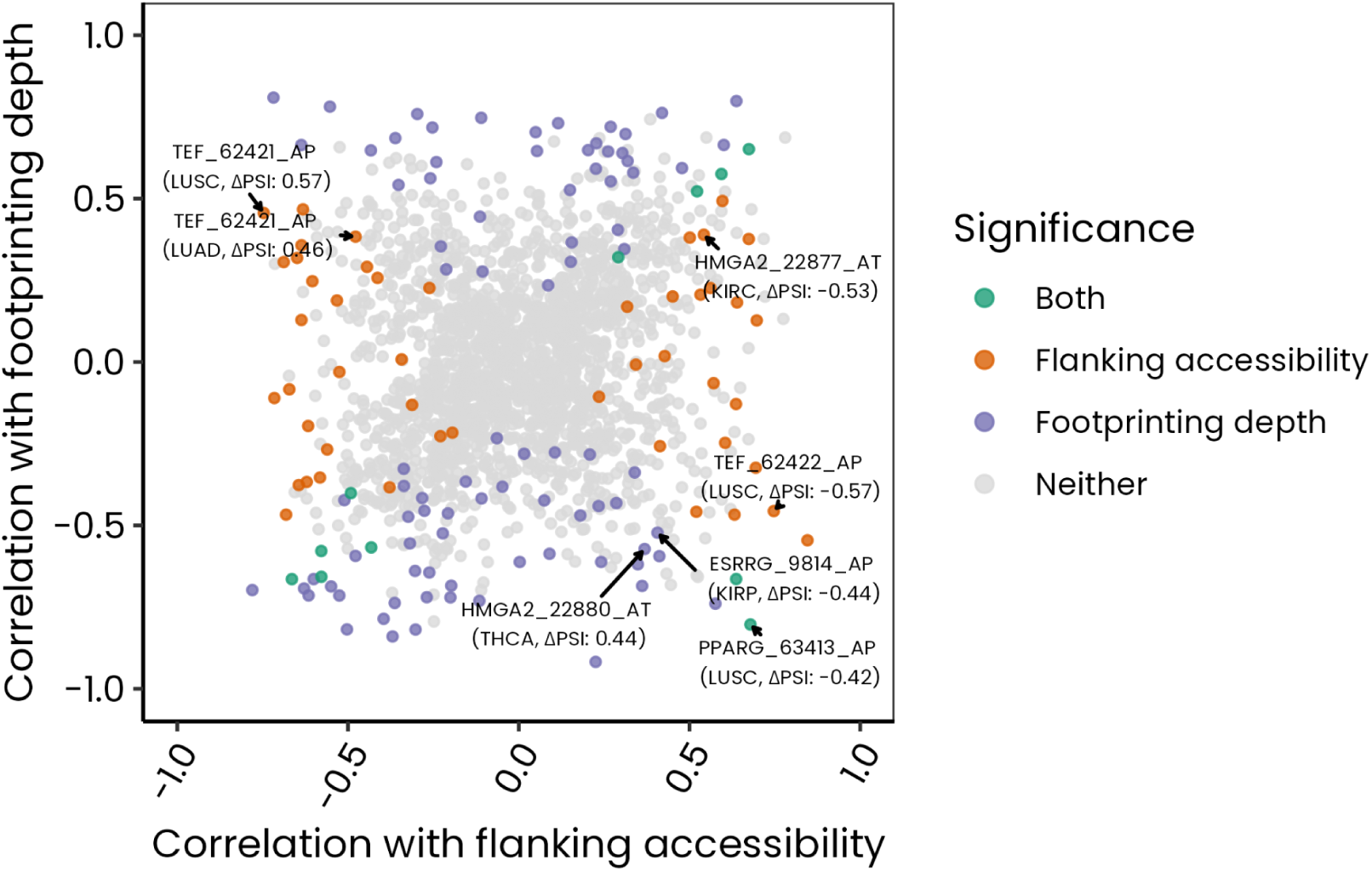
PTSEs correlated with footprinting. Each point represents one of the 1 948 PTSE-cancer type combinations (Table 2). Note that one PTSE-cancer type combination can have multiple footprinting depth/flanking accessibility values because of the presence of multiple DNA motifs for the corresponding TF (Section 2.6). In this case, the maximum correlation value (for each of footprinting depth and flanking accessibility) was taken to generate this figure.

### 3.6 PTSEs include TFs showing cancer dependencies

Loss-of-function screens in cancer cell lines, e.g., using Clustered Regularly Interspaced Short Palindromic Repeats (CRISPR)[40], have identified genes essential for cancer cell line survival. Because many TFs have been associated with the sustained cancer cell state[12], we evaluated the extent to which the TFs within PTSEs show cancer dependency, using the CRISPR-Cas9 screening data from The Cancer Dependency Map (DepMap, Broad (2025). DepMap Public 25Q2. Dataset. depmap.org)[20] (Section 2.7). Given a gene and a cancer cell line, DepMap provides gene perturbation effects based on Chronos score. A Chronos score of 0 or above indicates that the gene is not essential for the cancer cell proliferation, while a Chronos score of below 0 indicates the gene is essential for the cancer cell line proliferation, with the more negative value indicating more essentiality.

We found 669 TFs covered by PTSEs across 12 cancer types that had cancer dependency data for at least one cancer cell line (Figure 7A). 646 of the 669 TFs showed a negative Chronos score in cell lines associated with at least one cancer type (Supplementary Table S7). 39 of these TFs showed high (Chronos score < −1) dependency in a cancer cell line (Figure 7B-M). An interesting example is the PTSE *NKX2-1_27289_ES* affecting Homeobox protein Nkx-2.1 (NKX2-1) in LUAD (mean ΔPSI of −0.3). NKX2-1 has two known isoforms[41], where the PTSE *NKX2-1_27289_ES* leads to a shorter protein isoform. Because the mean ΔPSI value of NKX2-1_27289_ES is negative, the shorter isoform NKX2-1 potentially shows higher prevalence in cancer samples than in normal samples. NKX2-1 is a lineage TF for lung[42]. Functional differences between the two isoforms have also been shown[43]. Because the minimum Chronos score of NKX2-1 is −1.4 (Supplementary Table S7), indicating its essentiality in cancer proliferation, perhaps it’s the higher prevalence of the shorter isoform that determines the cancer dependency of NKX2-1. Overall, this analysis identified TFs that are not only essential for cancer proliferation, but also showed significant differences between normal and paired cancer samples with respect to AS events. This could imply that cancer dependency of these TFs are reliant on the corresponding TF isoforms.

**Figure 7.**
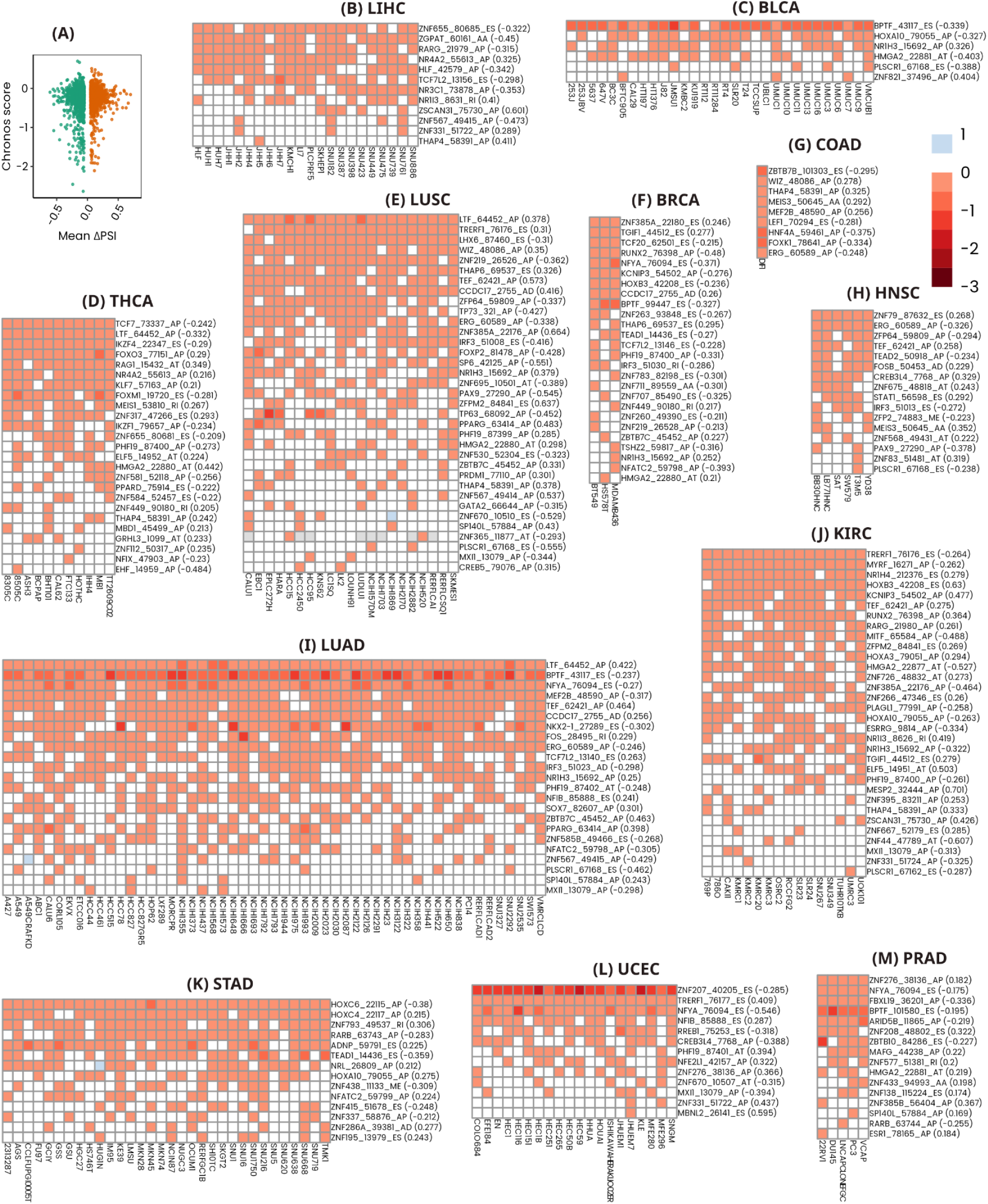
Cancer dependency of TFs involved in PTSEs. **(A)** Distribution of Chronos scores of all 2 505 TF-cancer type combinations (Section 2.7). Each point represents a TF-cancer type combination with x-axis representing the maximum ΔPSI value among all PTSEs affecting the TF and the y-axis representing the minimum Chronos score of the TF among all cell lines of the cancer type. **(B)-(M)** Chronos scores of individual cancer types, with x-axis showing the cell line and the y-axis showing the top 1% of PTSEs with maximum absolute mean ΔPSI value (denoted within brackets).

## 4 Discussion

Many AS events showing significant differences between paired normal and cancer samples were found across 14 cancer types, affecting nearly 50% of known TFs with sufficient annotations. Several of these TFs were found to be common in multiple cancer types, and many others were uniquely affected in a cancer type. These results highlight a pancancer significance of aberrant TF splicing in cancer. We found evidence of the impact of cancer-specific AS events on the DNA-binding and gene regulatory (activating/repressing) abilities of TFs. Further evidence for the functional impact of cancer-specific AS events on TFs were found by exploring whether the affected TFs were essential for cancer cell line proliferation. Overall, these findings could be used by future research to understand AS-driven cancer molecular mechanisms, which could guide identifications of novel therapeutic targets.

Different algorithms to detect and quantify AS events have been proposed in the literature, e.g., SpliceSeq[44], Sp1Adder[45], and Cufflinks[46]. All of the findings in this work are based on the TCGA SpliceSeq database, which used the SpliceSeq algorithm to quantify AS events. Future work could evaluate whether these findings are robust across different algorithms.

Gene regulatory associations based on gene level expressions of TFs have provided interesting insights in cancer[47]. Our results show that there is a large-scale potential functional impact of cancer-specific AS events on TFs, which could perturb gene regulatory associations. Future inference of gene regulatory associations in cancer could benefit from considering expressions of TFs not just on the gene level but also on the isoform level.

## Author contributions

KN and JB conceived the study. KN performed all of the work. KN, OT, and JB analyzed the results. JB provided the computational infrastructure. KN, OT, and JB wrote the manuscript.

